# Low rank approximation of difference between correlation matrices by using inner product

**DOI:** 10.1101/2021.02.23.432533

**Authors:** Kensuke Tanioka, Satoru Hiwa

**Affiliations:** Department of Biomedical Sciences and Informatics Doshisha University, Kyoto, Japan

## Abstract

**Introduction:** In the domain of functional magnetic resonance imaging (fMRI) data analysis, given two correlation matrices between regions of interest (ROIs) for the same subject, it is important to reveal relatively large differences to ensure accurate interpretations. However, clustering results based only on difference tend to be unsatisfactory, and interpreting features is difficult because the difference suffers from noise. Therefore, to overcome these problems, we propose a new approach for dimensional reduction clustering.

**Methods:** Our proposed dimensional reduction clustering approach consists of low rank approximation and a clustering algorithm. The low rank matrix, which reflects the difference, is estimated from the inner product of the difference matrix, not only the difference. In addition, the low rank matrix is calculated based on the majorize-minimization (MM) algorithm such that the difference is bounded from 1 to 1. For the clustering process, ordinal *k*-means is applied to the estimated low rank matrix, which emphasizes the clustering structure.

**Results:** Numerical simulations show that, compared with other approaches that are based only on difference, the proposed method provides superior performance in recovering the true clustering structure. Moreover, as demonstrated through a real data example of brain activity while performing a working memory task measured by fMRI, the proposed method can visually provide interpretable community structures consisted of well-known brain functional networks which can be associated with human working memory system.

**Conclusions:** The proposed dimensional reduction clustering approach is a very useful tool for revealing and interpreting the differences between correlation matrices, even if the true difference tends to be relatively small.

## Introduction

Currently, the neural basis of the human cognitive system is being studied using non-invasive neuroimaging techniques such as functional magnetic resonance imaging (fMRI), electroencephalography (EEG), and functional near-infrared spectroscopy (fNIRS) [1, 2, 3, 4]. In particular, to investigate the complex and distinctive functional network structure of the human brain and its nervous system [5, 6], functional connectivity analysis, which examines the temporal synchronization between brain regions (e.g [7]), is gaining popularity in this field. Functional connectivity between specific regions of interest (ROIs) is usually compared among various subjects or experimental conditions. Additionally, the patterns of functional connections are often analyzed in terms of network structures such as community structure and network centrality. For example, recent studies have revealed that the community structures of functional connectivity networks differ not only between schizophrenic individuals and healthy controls[8], but also during performance of different cognitive tasks[9].

Here, we focus on situations in which correlation matrices between ROIs are calculated for each subject in two different conditions. In such situations, it is important to reveal subnetworks of ROIs such that the difference between conditions is relatively large. However, it is difficult to interpret features of distinctive clusters because the range of correlations is bounded, i.e., [−1, 1] and the difference is affected by observational error. One very useful approach for addressing this problem is low rank approximation for the difference matrix. For example, low rank approximation combined with a heatmap, which is a data visualization tool, allows one to visually interpret the relations between ROIs. However, such estimated low rank matrices occasionally lose clustering structure information because the original difference matrix is affected by noise. To overcome this problem, we focus on the inner product matrix of the difference matrix. Even if the original difference matrix includes noise, the inner product matrix can emphasize the clustering structure of variables. Such an approach, which uses inner products for clustering, was proposed by [10] for processing high-dimensional data. To estimate low rank matrices, singular value decomposition is very popular. However, the range of the estimated low rank matrix is not bounded, while that of the correlation is bounded from −1 to 1.

In this paper, we propose a new dimensional reduction clustering approach for the inner product of the difference between two correlation matrices. Our approach has two advantages. First, the clustering results are superior to those from methods that rely on only the difference. Second, the range of the estimated low rank matrix is bounded from −1 to 1, and thus it is easy to interpret the relations. This approach consists of two steps. First, a low rank matrix is estimated such that each cluster is discriminated. Second, a clustering structure is calculated from the low rank matrix. A visualization of the proposed approach is shown in Figure 1. To address the problem of estimating a correlation matrix from a non-correlation matrix, various methods have been proposed[11, 12, 13, 14, 15]. In the proposed approach, we need to estimate a low rank correlation matrix, which indicates the difference between correlations, from the non-correlation matrix. To solve the nearest low rank correlation matrix problem [16, 17, 18, 19], we adopted the idea of majorization [16, 17]. We adopted and implemented this idea because it eases the estimation of low rank correlation matrices by using the majorization function[20, 21].

**Figure 1:**
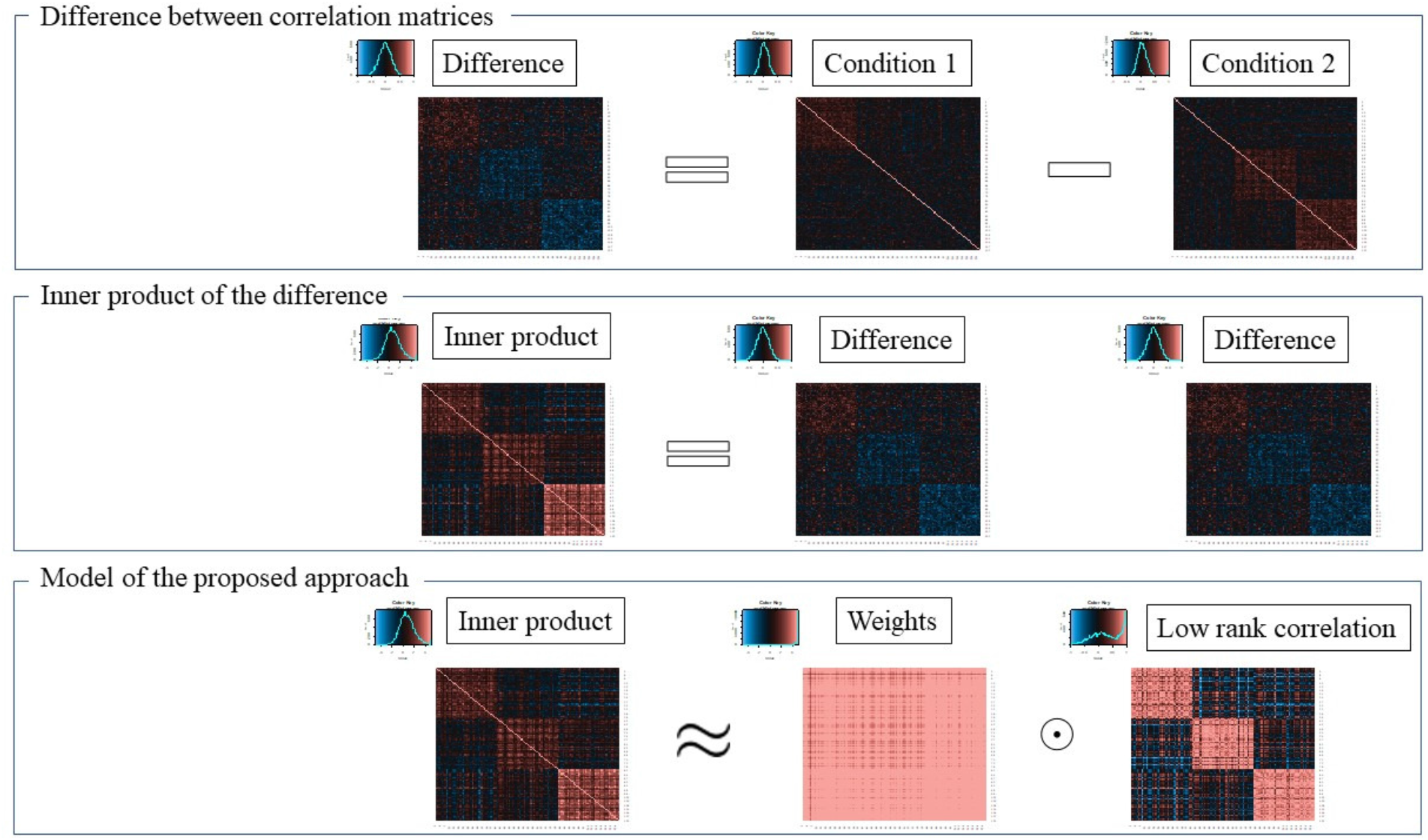
Image of the proposed approach

The remainder of this paper is organized as follows. In Methods, we discuss the reason for using the inner product of the difference between correlation matrices, and explain why this is better than using only the difference. In addition, a proposed model and corresponding objective function are introduced. To estimate the parameters, an algorithm based on the derived majorization function is provided. Next, the simulation design of our numerical study and the fMRI data for a mental arithmetic task are described. Then, the simulation results and real fMRI data for a mental arithmetic task are discussed. Finally, we offer our concluding remarks on the proposed method.

## Methods

In this section, we explain the proposed method. First, before introducing the optimization problem, we explain our reason for using the inner product of the difference between correlation matrices, instead of using only the difference. Next, the model of the proposed method is introduced, and the optimization problem of the proposed method is shown. Finally, the simulation design of our numerical study and the fMRI data for a mental arithmetic task are explained.

### Model of the proposed method

Here, the model of the proposed method is explained. Let 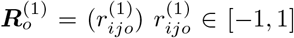, (*i, j* = 1, 2, · · · , *p*; *o* = 1, 2, · · · , *n*) and 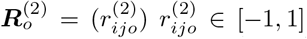, (*i, j* = 1, 2*, · · · , p*; *o* = 1, 2*, · · · , n*) be the correlation matrices between variables under condition 1 and condition 2, respectively, where *n* is the number of subjects and *p* is the number of variables. From 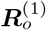 and 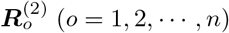, the difference is calculated as follows:

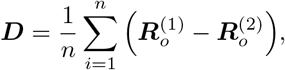

where ***D*** = (***d***_1_, ***d***_2_, , ***d**_p_*)^*T*^ = (*d_ij_*)*, d_ij_* ∈ [−2, 2] (*i, j* = 1, 2, · · · , *p*). The clustering structure of ***D*** is assumed to be in the following form:

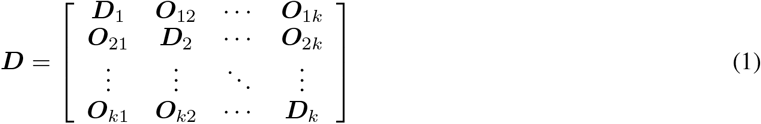

where ***D**_ℓ_* (*ℓ* = 1, 2, · · · , *k*) is a block matrix such that the absolute values of elements tend to be higher than those of elements of ***O**_ℓs_* (*ℓ* ≠ *s*). That is, ***D*** is assumed to be changed to Eq. (1) through permutation of these rows and columns.

However, even if ***D*** has such a clustering structure, the structure is masked because *d_ij_* are observed with noise and the corresponding correlations are bounded from −1 to 1. Therefore, it is difficult to capture the clustering structure by using low rank approximation based on only ***D***. To overcome this problem, the inner product of ***D*** is focused on because the clustering structure of the inner product is emphasized, rather than using only ***D***. Here, the inner product of ***D*** is defined as follows:

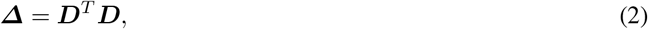

where elements of ***Δ*** are described as *δ_ij_* ∈ ℝ (*i, j* = 1, 2, · · · , *p*). We focus on the inner product and construct the model based on low rank approximation from *δ_ij_*. First, from the property of the inner product, for an arbitrary *δ_ij_*, there exists *θ_ij_* (0 < *θ_ij_* ≤ 2*π*) such that

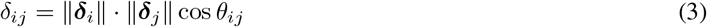

where ***δ**_j_* = (*δ*_1*j*_, *δ*_2*j*_, · · · , *δ*_*pj*_)^*T*^ (*j* = 1, 2, · · · , *p*) and || · || is the Euclidean norm. Here, cos *θ_ij_* can be considered as correlation, and the matrix representation is

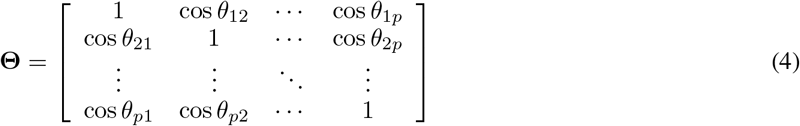

From Eq.(3) and Eq.(4), Eq.(2) is described as follows:

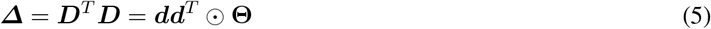

where ***d*** = (||***d***_1_||, ||***d***_2_||, · · · , ||***d**_p_*||)^*T*^, and ☉ is a Hadamard product.

Next, from Eq. (5), we construct following model:

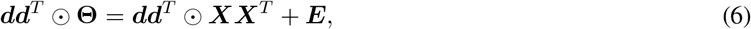

where ***X*** = (*x_iq_*) *x_iq_* ∈ ℝ(*i* = 1, 2, · · · , *p*; *q* = 1, 2, · · · , *d*) is a coordinate matrix with rank *d* (*d* ≤ *p*), *d* is determined by the researchers, and ***E*** = (*e_ij_*) *e_ij_* ∈ ℝ (*i, j* = 1, 2, · · · , *n*) is an error matrix. In addition, the following constraint is added to ***X***:

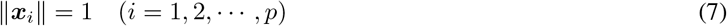

where ***x**_i_* = (*x*_*i*1_, *x*_*i*2_, · · · , *x_id_*)^*T*^ . From the constraint of Eq. (7), 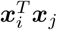 becomes the correlation coefficient between *i* and *j* with rank *d* [16, 17], and ***XX**^T^* also becomes a correlation matrix with rank *d*.

Again, the purpose of the proposed method is estimating a low rank matrix such that the clustering structure is emphasized. To achieve the purpose, ***X*** is estimated such that the sum of squares for ***E*** in Eq. (6) is minimized.

### Formulation of the proposed method

In this subsection, we show the proposed dimensional reduction clustering approach based on Eq.(6). The proposed method consists of two steps. First, a low rank correlation matrix, which indicates the difference between two correlation matrices, is estimated. Second, a traditional clustering algorithm such as *k*-means [22] is applied to ***X*** to obtain the clustering structure of variables. Although such two-step approaches have been proposed by [23, 24], these methods were proposed for multivariate data, not square matrices.

Next, the optimization problem of estimating a low rank correlation matrix is shown. Given ***Δ*** and *d*, the optimization problem is formulated as follows:

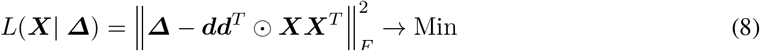

subject to

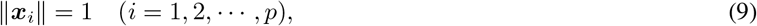

where || · ||_*F*_ indicates the Frobenius norm. In this study, to estimate ***X*** such as minimizing Eq. (8) with the constraint of Eq. (9), the majorize-minimization (MM) algorithm is used.

### Algorithm for estimating low rank correlation matrix based on MM algorithm

This subsection provides a detailed description of the MM algorithm for estimating ***X***. First, we derive the majorizing function of Eq. (8) in the same manner as [20, 21]. Next, the updated formula of ***x**_i_* is also derived based on the majorizing function. Then, based on the majorizing function, the problem is converted into a linear problem. Therefore, the updated formula can be derived by the Lagrange multipliers method with the constraint Eq. (9).

Now, the objective function Eq.(8) can be described as follows:

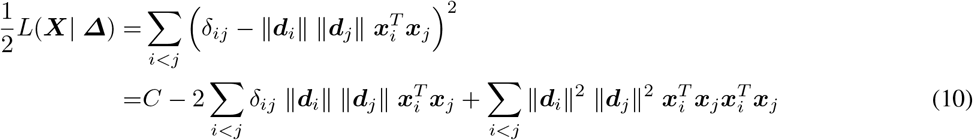

The third term of Eq. (10) corresponding to *i* is described as follows:

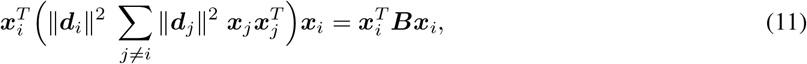

where 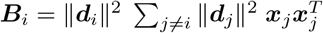. Let *λ_i_* be the maximum eigenvalue of ***B**_i_*. For any ***x*** ∈ ℝ^*d*^ with the constraint ||***x***|| = 1, the following inequality is satisfied:

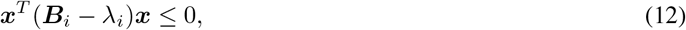

where ***I**_d_* is an identity matrix of size *d*. That is, ***B**_i_* − *λ_i_* is negative semi-definite. Let ***x**_i_* ∈ ℝ^*d*^ and ***z**_i_* ∈ ℝ^*d*^ be coordinate vectors of subject *i* corresponding to the current step and previous step, respectively. From Eq. (12), the following inequality is satisfied;

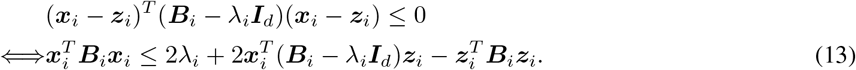

If ***x**_i_* = ***z**_i_*, Eq. (13) becomes an equality equation. Eq. (13) is substituted into the third term of Eq.(10) and we have

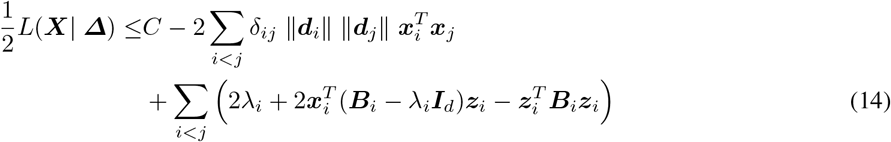

From (14), the updated formula of ***x**_i_* is derived as follows:

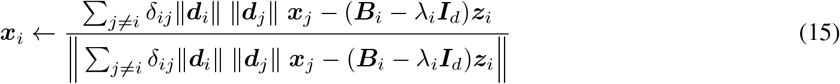

For the algorithm based on Eq. (15), see **Algorithm 1**.

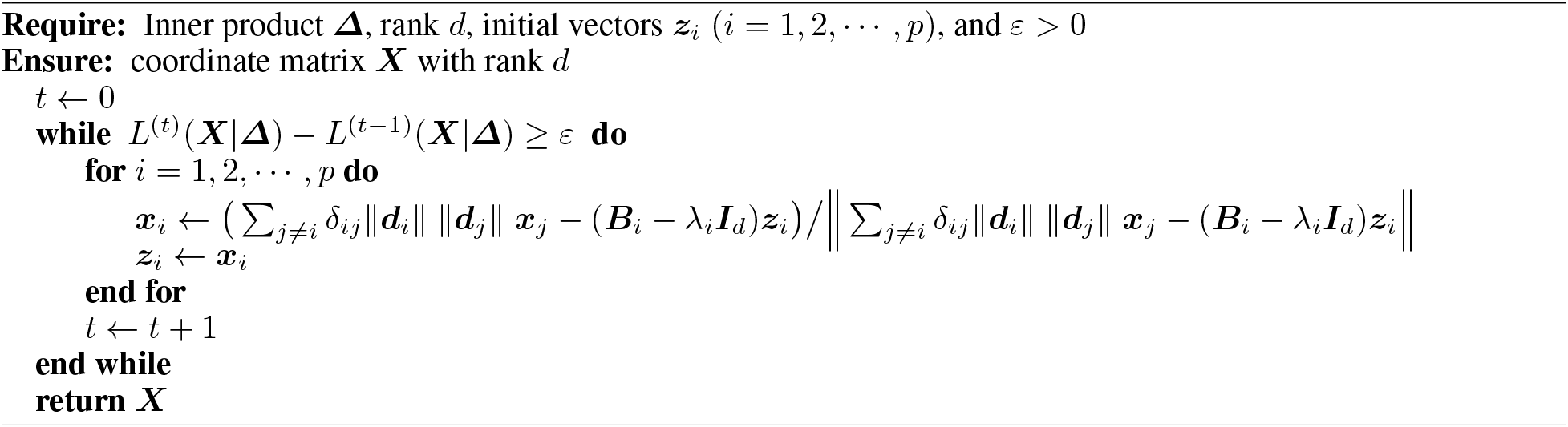
Algorithm 1. Estimating correlation matrix with rank *d*

Finally, to detect the clustering structure of variables, *k*-means is applied to ***X***. From results combined with a heatmap, we can interpret the clustering structures visually.

### Simulation study

In this subsection, the superiority of the proposed method is shown via the results of a numerical simulation. In particular, the recovery of clustering results is evaluated in this simulation.

We then reveal the simulation design. To evaluate the clustering results, artificial data with a true clustering structure are generated, and a correlation matrix between variables is then calculated from the data. Next, dimensional reduction clustering approaches are applied to the difference between two correlation matrices, and clustering results are obtained. Finally, an adjusted Rand index (ARI) [25] between the true clustering structure and the estimated clustering structure is calculated, and the effectiveness of the proposed method is compared with that of existing approaches.

Next, we explain the method for generating the artificial data. Multivariate data representing condition 1 and condition 2 are generated as follows:

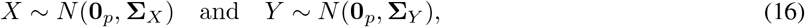

where *X* and *Y* are random vectors of conditions 1 and 2, respectively, **0**_*p*_ is a vector with length *p*, whose elements are all zero, and **Σ**_*X*_ ∈ [−1, 1]^*p×p*^ and **Σ**_*X*_ ∈ [−1, 1]^*p×p*^ are true correlation matrices of condition 1 and condition 2, respectively. Here, in this simulation, *p* is set to 120. From Eq.(16), *p* dimensional vectors are generated 60 times for each condition, and sample correlation matrices of condition 1 and condition 2 are calculated as ***R***(*X*) ∈ [−1, 1]^*p×p*^ and ***R***(*Y*) ∈ [−1, 1]^*p×p*^, respectively. Then, the input data are calculated as ***D*** = ***R***(*X*) − ***R***(*Y*).

In this simulate, four factors are utilized; a summary of the simulation is shown in Table 1. As a result, there are 4 × 3 × 3 × 2 = 72 patterns in this simulation. For each pattern, the corresponding artificial data are generated 100 times and evaluated using the ARI. Next, the descriptions of the factors are listed.

**Table 1:** Summary of the simulation design

| Names of factors | levels | descriptions |
| --- | --- | --- |
| Methods | 4 | proposal, Method2, Method3, and Method4 |
| Rank | 3 | Rank = 2, 3 and 4 |
| The number of clusters | 3 | $k = 2, 3$, and $4$ |
| The difference between true correlation | 2 | 0.1, and 0.2 |

### Factor 1: Methods

In this factor, we evaluate 4 methods. The first method is the proposed approach based on the inner product of ***D***. For the second method, the approach based on difference, not the inner product model, is adopted. The second method consists of two steps, as does the proposed method. First, ***D*** is decomposed as ***D*** = ***B**^T^ **B*** by using Cholesky decomposition, where ***B*** = (***b***_1_*, **b***_2_, · · · , ***b**_p_*) ***b**_i_* ∈ ℝ^*q*^ (*i* = 1, 2, · · · , *p*; *q* ≤ *p*). Let ***b***^†^ = (||***b***_1_||, ||***b***_2_||, · · · , ||***b**_p_*||)^*T*^; therefore, there exists cos *ϕ_ij_* such that *d_ij_* = ||***b**_i_|| ||**b**_j_*|| cos *ϕ_ij_* (0 ≤ *ϕ_ij_* < 2*π*). From the decomposition, the optimization problem of the second method is formulated as follows:

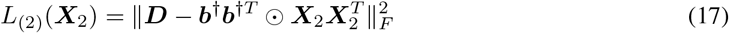

subject to

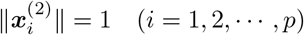

where *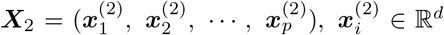*. Parameters of Eq.(17) can be estimated in the same manner as in the proposed method. Subsequently, *k*-means is applied to the estimated ***X***_2_. The third method also consists of two steps. Eigenvalue decomposition is applied to ***D***, and the low rank matrix corresponding to *λ*_1_*, λ*_2_, · · · , *λ_d_* is estimated, where *λ*_1_ ≥ *λ*_2_ ≥ · · · ≥ *λ_q_* (*q* ≤ *p*) are eigenvalues of ***D***. Afterward, *k*-means is applied to the estimated low rank matrix. The fourth method is similar to the third method; the only difference from the third method is that eigenvalue decomposition is applied to the inner product matrix of ***D***, not only to ***D***.

Both the third and fourth methods provide us with low rank matrices and clustering results. However, from these results, it is difficult to interpret the degree of the relation because these estimated values are not bounded. On the other hand, both the first and second methods allow us to interpret the results easily, as these estimated values are bounded from 1 to 1.

### Factor 2: Rank

In the four factor 1 methods, the rank must be determined. In this simulation, ranks are set as 2, 3, and 4.

### Factor 3: The number of clusters

All methods in factor 1 adopt *k*-means clustering. In the simulation, we assume a situation in which the number of true clusters is known beforehand. The number of clusters is set as *k* = 2, 3, , and 4.

### Factor 4: The difference between true correlations

At first, the generation of true clustering structures is determined by *k* = 2, 3, and 4. The true clustering structure is dependent on **Σ**_*X*_ and **Σ**_*Y*_ . When *k* = 2, **Σ**_*X*_ and **Σ**_*Y*_ are

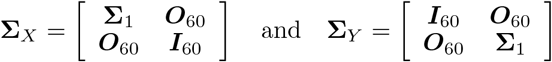

where ***O***_60_ ∈ {0}^60×60^, ***I***_60_ is a 60 × 60 identity matrix, and 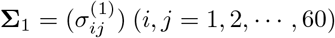 and 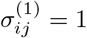 if *i* = *j*.

Again, there are two levels in factor 4. The first level and second level are 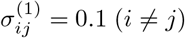 and 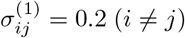, respectively.

For the case of *k* = 3,

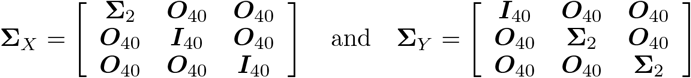

where **Σ**_2_ ∈ {0, 0.1, 0.2, 1}^40×40^ is defined in the same manner as **Σ**_1_. Finally, the case of *k* = 4:

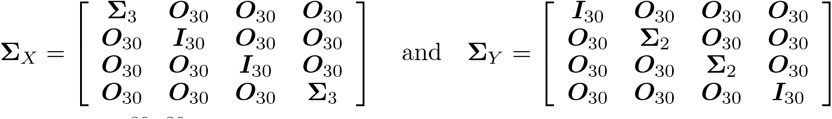

where **Σ**_3_ ∈ {0, 0.1, 0.2, 1}^30×30^ is defined in the same manner as both **Σ**_1_ and **Σ**_2_.

From the above **Σ**_*X*_ and **Σ**_*Y*_, a true clustering structure is described. In the cases of *k* = 2, 3, and 4, the cluster size of each is the same. That is, for *k* = 2, 3, and 4, the cluster sizes are 60, 40, and 30, respectively.

### fMRI data for mental arithmetic task

Among the cognitive functions, working memory (WM) is important for engaging in everyday tasks such as conversation or reading books. WM is the system for keeping the necessary information temporally and for processing the information [26, 27]. In studies on cognitive function, the neural basis of the WM has been investigated using fMRI. In current research, the brain regions and functional connectivity networks associated with the WM system are being revealed. Therefore, to validate the effectiveness of the proposed method, we apply the proposed method to fMRI data measured during the WM task.

### Participants

Thirty-two healthy adults (20 males and 12 females; mean age, 22.0 ± 1.2 years; 31 right-handed and 1 left-handed) participated in this experiment. This study was approved by the Research Ethics Committee of Doshisha University (approval code: 1331). Written informed consent was obtained from each participant.

### Experimental design

The participants were asked to perform a mental arithmetic task in an fMRI scanner. The experimental design is shown in Fig . 2. In this experiment, a block design was adopted such that the rest periods and task periods were conducted alternately. Participants were instructed to watch a fixation point during rest, and to press buttons to answer true or false in response to a presented numerical formula during a task. They could proceed to the next formula at their own pace by selecting an answer to the current formula. They performed two types of mental arithmetic tasks, each having a different level of difficulty: 1) low-WM load (Low-WM) tasks consisted of addition of one-digit numbers, and 2) high-WM load (High-WM) tasks consisted of arithmetic operations on real numbers with three digits. Each of the two task types was performed three times (each participant performed a total of six tasks in the experiment) and the task order was randomized. The durations of the first rest block, second rest block, and task blocks were 40s, 36s, and 30[s], respectively.

**Figure 2:**
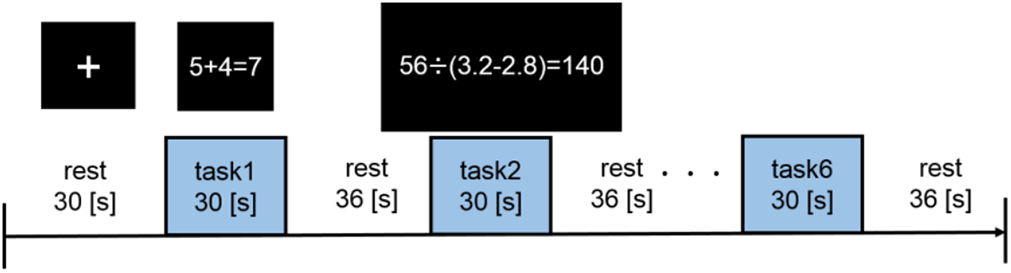
Experimental design

### Data acquisition

All MRI scans were performed on a 1.5T Echelon Vega (Hitachi, Ltd). Functional images were acquired using a gradient-echo echo-planar imaging sequence (TR=3000ms, TE=40ms, flip angle = 90°, field of view = 240 × 240, matrix size = 64 × 64 pixel, thickness = 5.0mm, slice number = 20). Structural images were acquired using an Rf-spoiled steady state gradient echo sequence (TR=9.4ms, TE=4.0ms, flip angle = 8°, field of view = 256 × 256 mm, matrix size 256 × 256 pixel, thickness = 1.0 mm, slice number =194). The stimuli (the fixation point and arithmetic formula) were presented and synchronized with fMRI data acquisition using presentation software (Neurobehavioral System Inc., Albany CA), and participant responses were acquired by the fORP932 Subject Response Package (Cambridge Research Systems Ltd., London UK).

### Data preprocessing

The first six scans were excluded from the analysis in order to eliminate the nonequilibrium effects of magnetization. The functional images were preprocessed using SPM12 software (Wellcome Department of Cognitive Neurology, London UK) [2]. They were realigned to correct for head movements, and subsequently slice-timing corrected; anatomical images were then coregistered to the mean of the functional images. They were also spatially normalized into Montreal Neurological Institute (MNI) space and smoothed using a Gaussian filter (8mm full width-half maximum).

### Functional connectivity analysis: Deriving the correlation matrices

For the functional connectivity analysis, the functional images preprocessed using SPM12 were further processed using the CONN toolbox [28]. In detail, nuisance regression was performed using an anatomical component-based noise correction method (aCompCor) [29], in which the first 5 principal components of the signal from the white matter and cerebrospinal fluid masks, as well as the head-motion parameters, their first-order temporal derivatives, and the main task effect from the task block (modeled as a canonical hemodynamic-response-function-convolved response) and its first order derivatives, were regressed out to eliminate physiological noise and the potential confounding effects of task responses.

To calculate functional connectivity during tasks, the preprocessed images were parcellated into 116 regions, including 90 cerebrum regions and 26 cerebellum regions defined by automated anatomical labeling (AAL); the mean Blood-Oxygen-Level Dependent (BOLD) time course was then calculated for each region. Subsequently, Pearson correlations among BOLD time courses of 116 regions were calculated and then Fisher-z transformed. As a result, a 116 × 116 correlation matrix was constructed for each participant. In this study, we chose 90 cerebrum regions as the ROIs and used the corresponding 90 × 90 correlation matrices for the analysis because the WM system is considered to be associated with the cerebrum regions [30].

## Results

In this section, we describe the results of numerical simulations and of applying the proposed fMRI data method.

### Simulation results

In this subsection, we describe the results of numerical simulations.

Figure 3 shows results in terms of the number of clusters and ranks in the case of 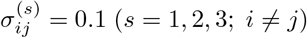. In this situation, we assume that the true difference is relatively small. Therefore, it is difficult to recover the true clustering structure. The vertical axes represent results of ARIs. Here, the methods are compared in terms of the medians and interquartile ranges (IQRs) of the ARIs. In Figure 3, the results of the second and third methods, which utilize only the difference and not the inner product, indicate lower performance than that of the proposed method. As the results in Figure 3 show, the model that utilizes the inner product tends to outperform models that focus only on the differences. In particular, the results of the proposed method tend to be superior to those of the other methods in the case of *d* ≥ *k*.

**Figure 3:**
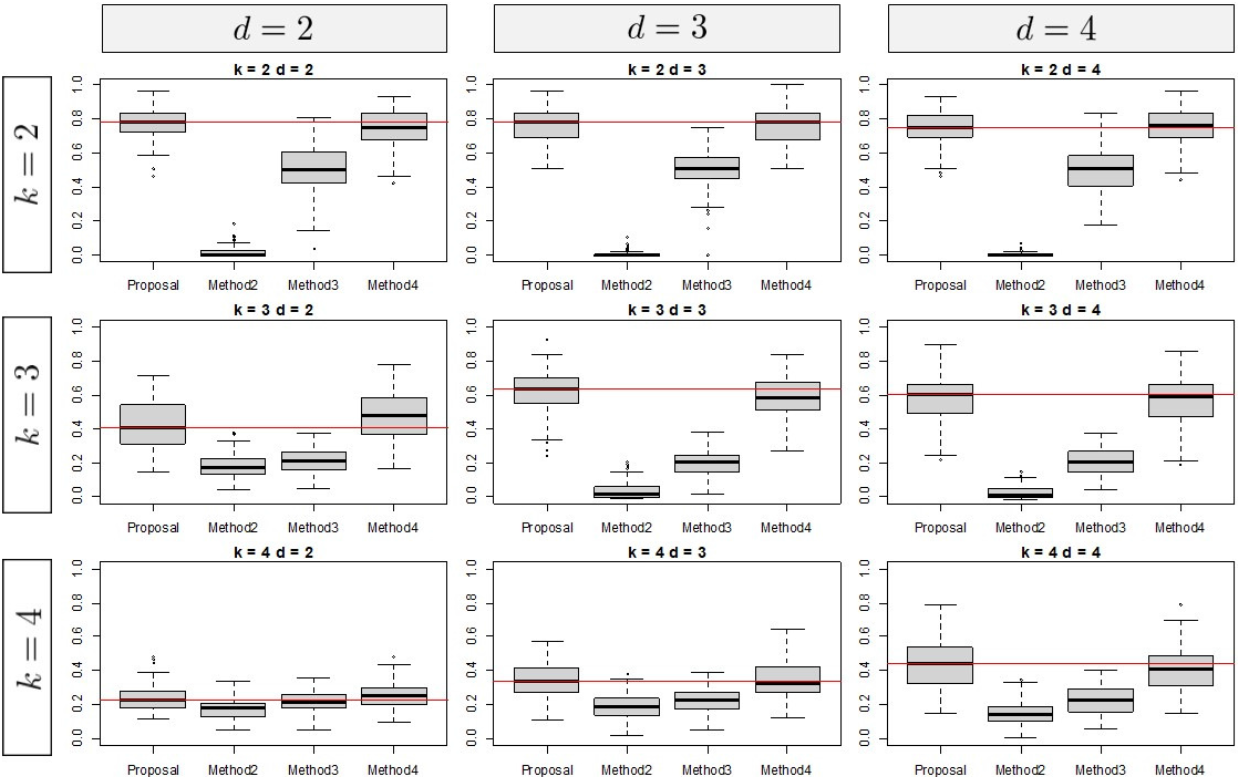
Simulation result of case 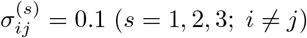

The results of 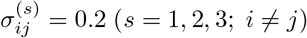 are shown in Figure 4. The tendency of these results is similar to those of 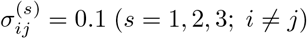, while the results corresponding to 0.2 tend to indicate higher performance than those corresponding to 0.1.

**Figure 4:**
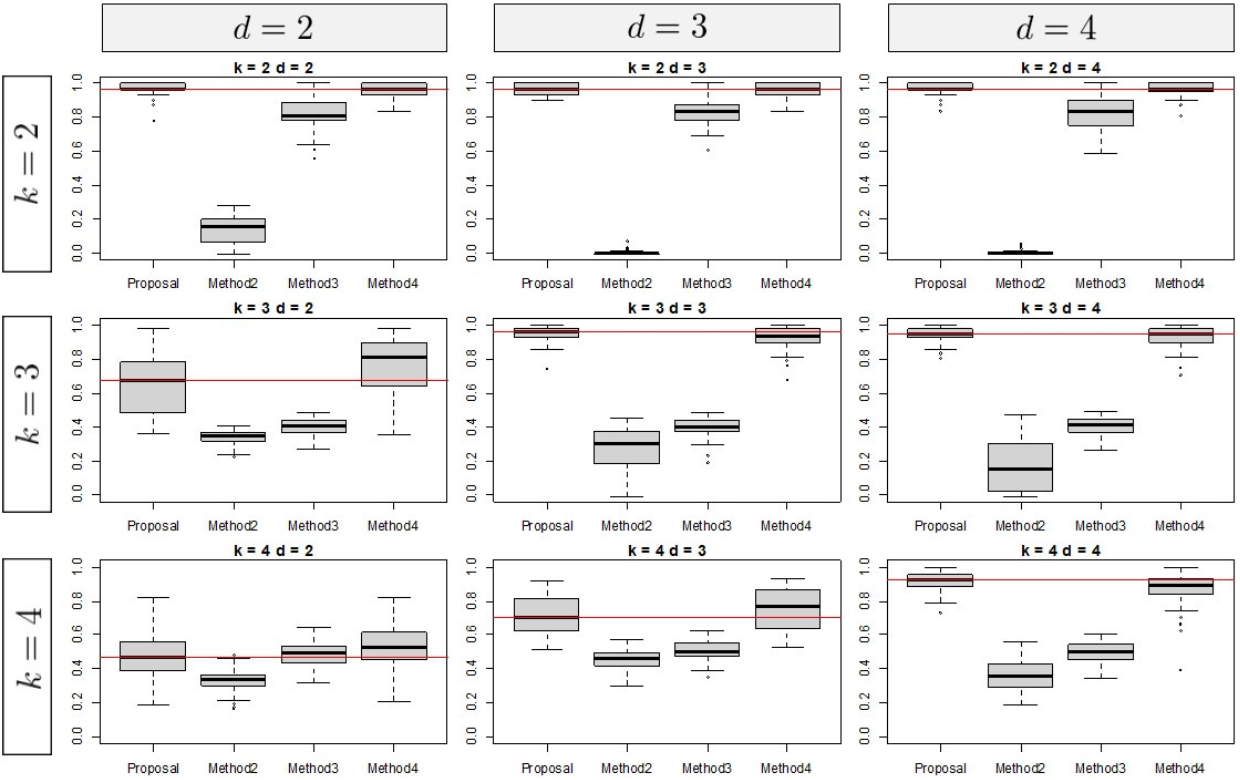
Simulation result of case 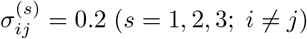

From the overall results, we note two specific observations. First, the number of clusters is relatively large, and performance tends to be relatively low irrespective of the type of method; this tendency is pointed out by [31]. Second, all methods depend on the selection of hyperparameters. Based on the simulation results, the tuning parameters should be set as *d* ≤ *k*.

### Results of fMRI data analysis

In this subsection, we show the results of applying the proposed method to the fMRI data. Concretely, the purpose of this example is detecting clustering structures where the difference between two experimental conditions, i.e., High-WM and Low-WM tasks, is emphasized. In addition, features of these estimated clusters are interpreted combined with knowledge of ROIs related to WM, including the task positive network (TPN), ventral attention network (VAN), salience network (SN), visual network (VN), and default mode network (DMN). The TPN consists of the fronto-parietal network (FPN), dorsal attention network (DAN), and cingulo-opercular network (CON). The FPN and DAN are related to executive function [32] and top down attention, respectively, and the CON is involved in exerting control over the contents of WM [33]. In addition, the VAN is associated with response to stimuli and bottom-up attention, and SN activation is said to be observed in situations in which it may be advisable to change behavior [34, 35]. The VN is literally associated with visual information processing, and the DMN relates to internally focused tasks such as autobiographical memory retrieval, envisioning the future, and conceiving the perspectives of others [36].

Next, we explain how to construct the difference between correlation matrices. For each subject, using a matrix of difference between correlation matrices of High-WM and Low-WM, we calculate one mean matrix of these differences. As described in the previous section, the input difference is a 90 × 90 matrix.

In the proposed method, rank and the number of clusters must be selected. For the determination of rank, we set the rank candidates as = 2, 3, 4, 5, 6, and select one among them. Generally, when the rank is larger, the values of the objective function tend to be lower.

The proposed method is then applied to the difference matrix using rank = 2, 3, 4, 5, 6, and the rate of change for values of the objective function is calculated as follows:

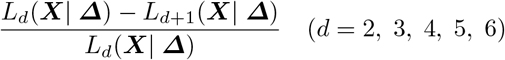

where *L_d_*(***X | Δ***) is the value of the objective function with rank *d*. From Figure 5, *d* = 3 is selected because the change of values for the objective function tend not to change even if the rank is larger than 3 among the candidates. For the number of clusters, silhouettes coefficient [37] is used; the number of clusters that provides the highest silhouettes coefficient is *k* = 3. See Figure 6. The left side of Figure 6 shows the results of applying the proposed method. Through the estimated low rank correlation matrix, we confirm that the clustering structure is emphasized, and that the relations among ROIs belonging to the same cluster are estimated higher. The right-hand side of Figure 6 shows the original difference matrix, in which both rows and columns are permutated by the estimated clusters. The right-hand side of Figure 6 also shows that relations among ROIs belonging to cluster 1 tend to be higher under the difficult task than under the easy task, although those among ROIs belonging to cluster 2 or cluster 3 tend to be lower.

**Figure 5:**
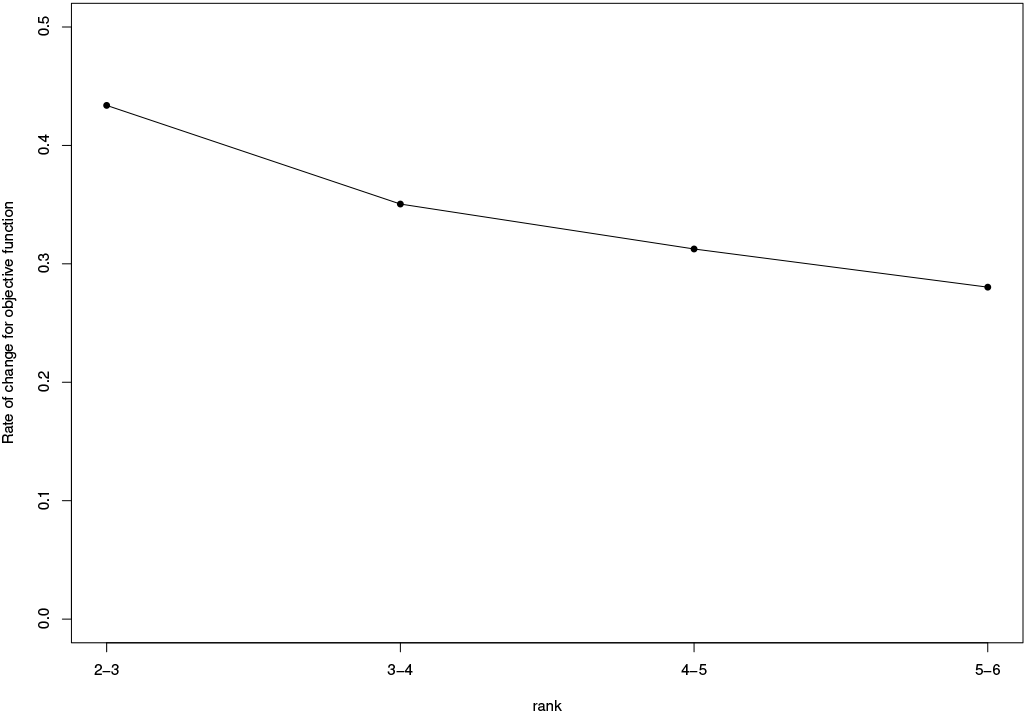
Change of rate for the objective function; vertical and horizontal axes indicates the rank and the ratio of values for the objective function, respectively.

**Figure 6:**
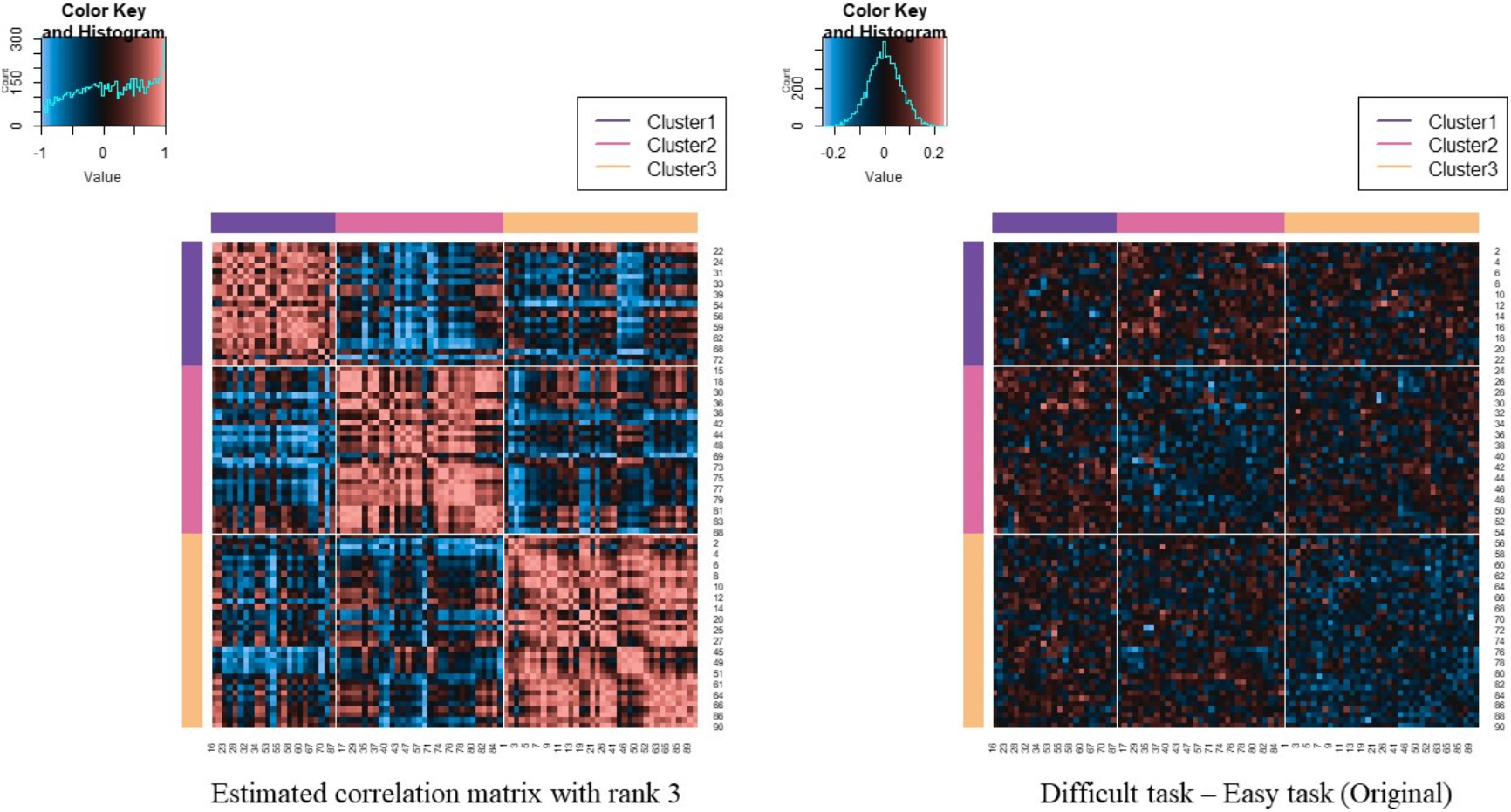
Estimated correlation matrix with *d* = 3 and the clustering result

Next, the features of the estimated cluster are interpreted combined with knowledge of WM. Figure 7 represents the difference matrix (High-WM - Low-WM) permutated by both of the estimated clusters and WM. We particularly focused on the differences related to the TPN and DMN. For the relations of ROIs belonging to the TPN, those belonging to cluster 1 tend to be higher than those belonging to cluster 2 or cluster 3. Therefore, in cluster 1, correlation among ROIs related to the TPN tended to be active under the condition of High-WM.

**Figure 7:**
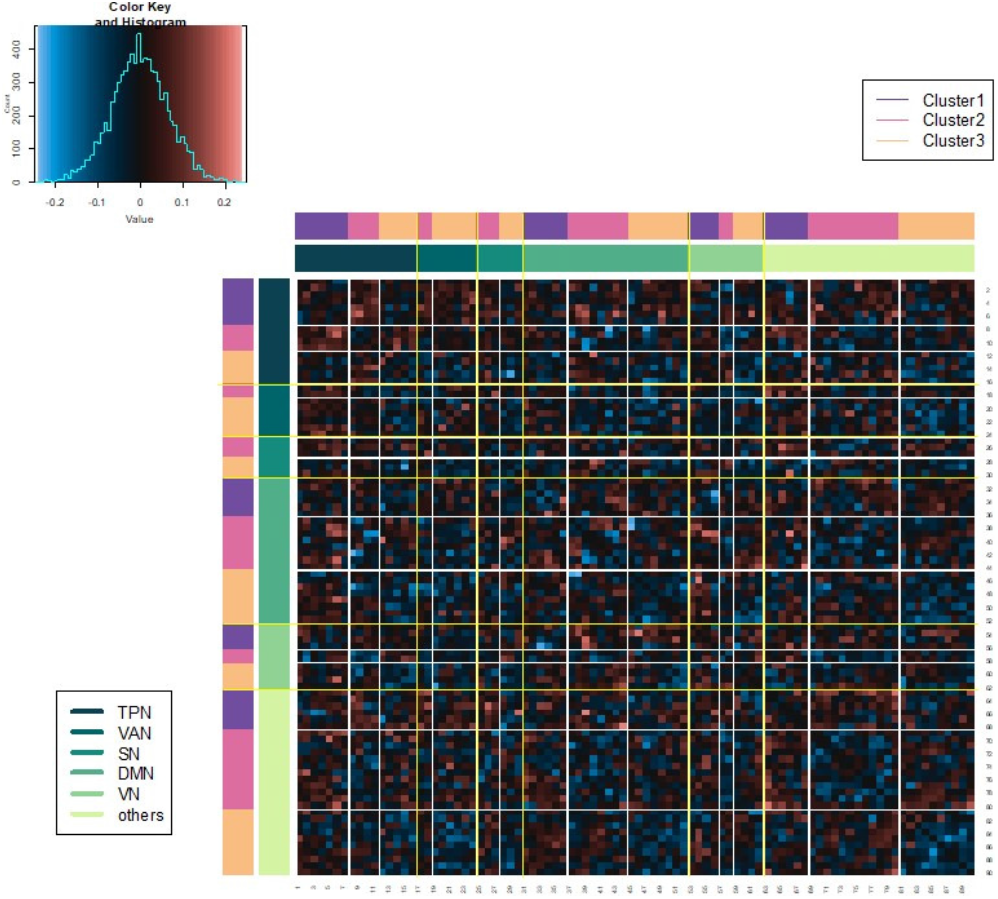
Difference matrix (High-WM - Low-WM) permutated by the estimated clusters and WM

To confirm the features of clusters, Figure 8 is shown. In Figure 8, each boxplot represents the differences between ROIs within WM for each cluster. When correlations within WM are active in the High-WM condition, the median tends to be greater than 0. On the other hand, the median tends to be lower than 0 when those are active in the Low-WM condition. First, the features of cluster 1 are interpreted. The values corresponding to the TPN in cluster 1 tend to be higher than those in cluster 2 and cluster 3. Again, the TPN includes ROIs within the FPN, DAN, and CON. The connectivities within the FPN tend to be higher in High-WM [38], and the DAN is related to top-down attentional control [39]. In addition, there are no ROIs belonging to the VAN or SN as features of cluster 1, where the VAN is related to bottom-up attentional processing. Therefore, cluster 1 is interpreted as ROIs related to top-down attentional control. In cluster 2, there are no ROIs belonging to either the FPN or DAN, and the TPN includes only the CON. The values for the SN in cluster 2 tend to be higher than those in cluster 1 and cluster 3; the SN facilitates accessing attention and working memory resources. In addition, the differences corresponding to the DMN tend to scatter around 0 symmetrically. From these features, cluster 2 is interpreted as ROIs related to the SN. In cluster 3, those within the TPN are lower compared to those within the other clusters. In addition, those within the VAN tend to be higher than those in cluster 2, where there are no ROIs within the VAN. Therefore, cluster 3 is interpreted as ROIs related to bottom-up attentional processing.

**Figure 8:**
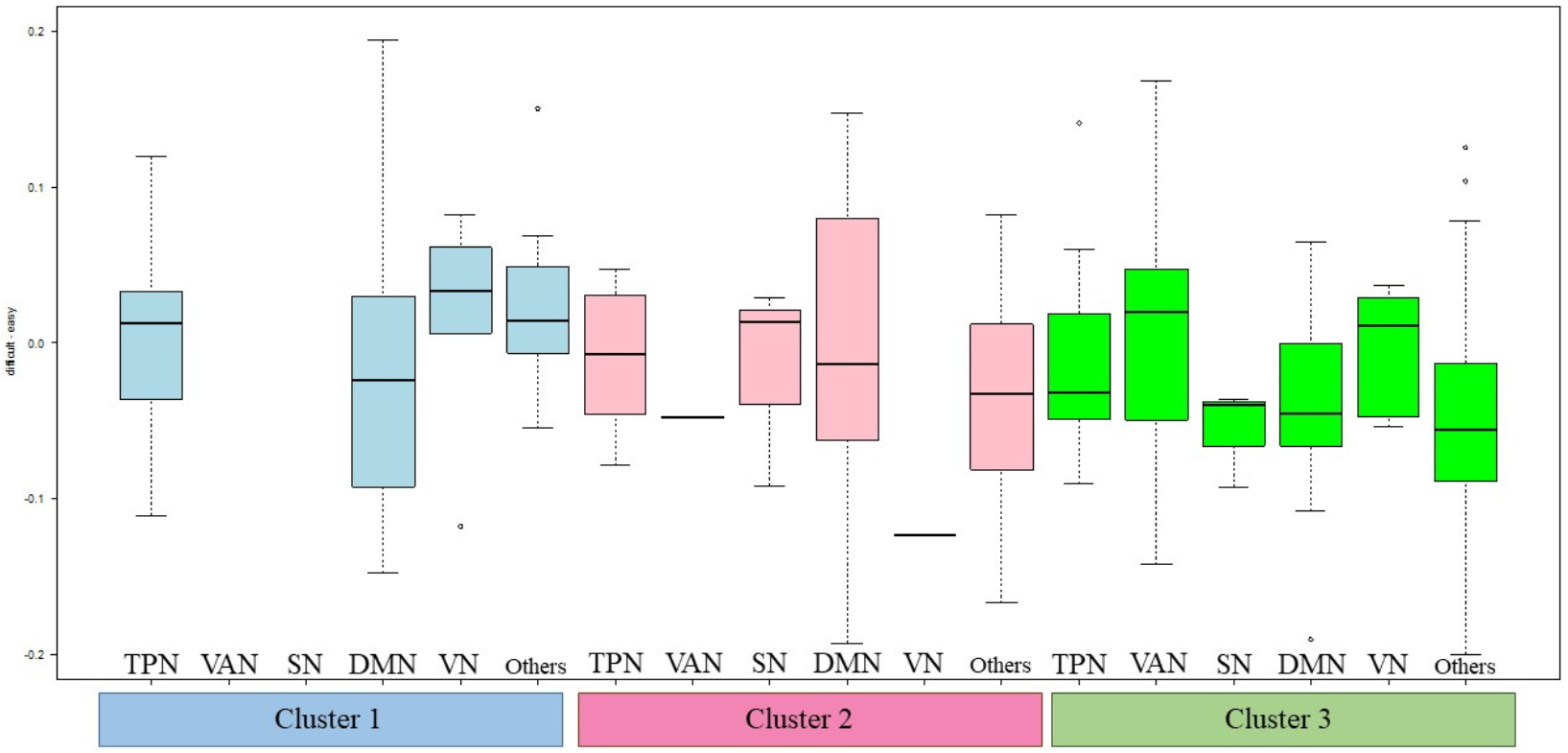
Boxplot of the difference (High-WM - Low-WM) by the estimated clusters and WM

Although the proposed method was applied to the fMRI data during mental arithmetic tasks as an explanatory analysis, these results suggest that our method can detect community structures consisting of well-known functional networks associated with the human working memory system.

## Conclusion and remarks

We proposed a new dimensional reduction clustering approach based on the difference between correlation matrices. The estimated low rank correlation represents the difference, and it is easy to interpret the relations as the estimated values are bounded from 1 to 1 and the clustering structure is emphasized. In addition, based on the idea of using the inner product of the difference, not only the difference, we demonstrated that the clustering recovery tends to be accurate. The effectiveness of the proposed method is demonstrated through numerical simulations of the recovery of a true clustering structure.

We show the results of applying the proposed method to real fMRI data related to WM. In fact, our proposed method provides the clusters, which are interpreted from the knowledge of WM.

## Acknowledgements

Not applicable.

## Abbreviations

AAL: automated anatomical labeling
ARI: adjusted rand index
aCompCor: an anatomical component-based noise correction method
BOLD: blood oxygen level dependent
CON: cingulo opercular network
DAN: dorsal attention network
DMN: default mode network
EEG: electroencephalography
fMRI: functional magnetic resonance imaging
fNIRS: functional near-infrared spectroscopy
FPN: fronto-parietal network
IQRs: interquantile ranges
MNI: Montreal Neurological Institute
ROIs: regions of interest
SN: salience network
VN: visual network
TPN: task positive network
VAN: ventral attention network
WM: Working Memory.

## Availability of data and materials

The artificial data used in numerical simulations is generated from the probability distribution.For the way of generation, see subsection Simulation study. The datasets used in this study are available from authors on reasonable request.

## Ethics approval and consent to participate

This study was approved by the Research Ethics Committee of Doshisha University (approval code: 1331). Informed consent were obtained for all subjects before enrolling the experiment.

## Consent for publication

Not applicable.

## Authors’ contributions

KT construct the proposed statistical method, conducted numerical simulations and applied the method to real fMRI data, drafted the initial manuscript. SH designed the experimental design, conducted the experiment, proposed the framework of fMRI data analysis, and reviewed the manuscript.

